# Chronic Proliferation and Glia Heterogeneity in CMT1A: Involvement of RUNX2-Positive Schwann Cells and Endoneurial Fibroblasts

**DOI:** 10.64898/2026.09.25.754314

**Authors:** Davide Visigalli, Mattia Camera, Davide Ceresa, Giovanna Capodivento, Daniela Marubbi, Angelo Schenone, Paolo Malatesta, Lucilla Nobbio

## Abstract

Charcot–Marie–Tooth type 1A (CMT1A) is the most common inherited peripheral neuropathy, caused by duplication of the peripheral myelin protein 22 (*PMP22*) gene and characterized by dysmyelination and defects in myelin maturation. Hypercellularity is a prominent feature of the disease, although its origin, persistence, and underlying mechanisms remain controversial. Here, we performed a longitudinal analysis of sciatic nerves from wild-type (WT) and CMT1A rats between 2 and 12 months of age.

We demonstrated that CMT1A nerves exhibit persistent cellular proliferation extending into adulthood. This phenotype is confirmed by bulk RNA-seq data showing sustained enrichment of proliferation-associated gene signatures. Cell-type signature analysis identifies overrepresentation of immature, neural crest-related, and epithelial-mesenchymal transitional (EMT) transcriptional programs in CMT1A nerves. Among the most upregulated genes, RUNX2 showed strong and prolonged expression at both transcript and protein levels. Consistently with the key role of RUNX2 in the activation of the injury response, CMT1A nerves also showed a marked enrichment in SOX10^+^/JUN^+^ cells, that only partially overlap with SOX10^+^/RUNX2^+^ cells, indicating a maladaptive activation of the injury-like Schwann cell (SC) program. Interestingly, we detected robust upregulation of SMOC2, an extracellular matrix protein selectively expressed by endoneurial fibroblasts, pointing to activation of non-canonical cell types and altered extracellular matrix remodeling.

Together, these data demonstrate that CMT1A peripheral nerves are characterized by chronic hypercellularity driven by persistent proliferation, EMT, and dysregulated injury transcriptional programs, collectively pointing at progressive and multicellular nerve remodeling as a relevant pathomechanism in the disease.

**Highlights:**

- CMT1A nerves exhibit persistent proliferation throughout adulthood
- CMT1A induces immature and injury prone transcriptional programs
- RUNX2 marks a distinct Schwann cell subpopulation in CMT1A nerves
- Endoneurial fibroblasts contribute to progressive nerve remodeling

## Introduction

Charcot-Marie-Tooth type 1A (CMT1A) is a hereditary peripheral neuropathy that typically manifests during childhood or adolescence. It represents the most common inherited neuropathy and the predominant subtype among all CMTs [1], [2]. It is caused by the genomic duplication of a region containing the *PMP22* gene, first identified more than thirty years ago [3], [4]. However, the downstream mechanisms through which *PMP22* duplication affects peripheral nerve physiology remain poorly understood.

Patients commonly present with gait abnormalities and distal motor impairment, as well as a characteristic and uniform slowing of conduction velocities [5]. These alterations arise early in life in CMT1A patients and are characterized by the loss of large myelinated axons, accompanied by a shift toward a more homogeneous population of smaller-caliber axons with a reduced g-ratio, resembling the structural features of immature peripheral nerves [6], [7], [8]. Thus, CMT1A is a dysmyelinating neuropathy, associated with defects in myelin and axonal maturation, and remarkable impairment of lipid metabolism [9], [10], [11].

In fact, CMT1A nerves are not only hypomyelinated but also undergo extensive remodelling involving nearly all structural components, including myelin, axons, extracellular matrix, and different cell types [12], [13]. In this context, cellular proliferation appears to play a fundamental role. Several studies have reported marked hypercellularity during the CMT1A nerve development, a finding reproduced in multiple CMT1A animal models and also observed in human nerves, where cell density may reach up to tenfold that of healthy controls [14], [15], [16], [17], [18], [19], [20]. Various interpretations have been proposed to explain this phenomenon, including onion bulb formation, low-grade inflammation, and proliferation within Schwann cell (SC) lineages. The cell types potentially involved include resident macrophages and several SC phenotypes as myelinating, non-myelinating, immature, and the supernumerary SCs associated with onion bulb structures [15], [16], [21], [22], [23], [24].

Cellular proliferation in CMT1A, however, remains a controversial issue. Some studies reported no evidence of proliferation in the disease, whereas others described proliferation accompanied by increased apoptosis, or proliferation limited to onion bulbs or just involving macrophages. Furthermore, *in vitro* studies have suggested that *PMP22* overexpression may act as a specific growth-arrest signal that inhibits cell proliferation and entry into the cell cycle [25]. In contrast, several reports documented increased proliferative activity in rodent models, although these studies were conducted within a timeframe limited to two months of age [15], [16], [23], [26].

To address these discrepancies concerning hypercellularity and proliferation, we performed a longitudinal analysis of adult sciatic nerves from 2 to 12 months of age in wild-type (WT) and CMT1A rats, thereby minimizing the contribution of physiological processes which occur during the very early stages of development to the observed proliferative activity. Our analyses revealed persistent cellular proliferation in CMT1A nerves. Transcriptomic profiling further demonstrated strong enrichment of epithelial-mesenchymal transitional (EMT) transcriptional programs, accompanied by sustained expression of RUNX2, a transcription factor recently involved in injury-mediated responses [27]. In addition, SMOC2, an extracellular matrix protein exclusively secreted from endoneurial fibroblasts, is increased in CMT1A sciatic nerves, supporting a possible alteration of fibroblasts physiology in CMT1A.

Overall, our findings indicate that CMT1A nerves undergo extensive and progressive tissue remodelling driven by persistent cellular proliferation, coupled with EMT and dysregulated injury programs, suggesting novel therapeutic targets to impact the disease.

## Material and Methods

### Animal model

The CMT rat, an animal model of CMT1A neuropathy originally developed in K.-A. Nave laboratory, was used for the experiments [28]. Heterozygous CMT1A and WT littermates of both sexes at postnatal day (P) 60, P180, and P365 were used for most experiments. We also performed experiments during the early stages of development (P5-P20). All experimental procedures were accomplished in accordance with the Italian Legislative Decree 26/2014 and the EU Directive 2010/63/EU for the care and use of laboratory animals. The protocol was approved by the Institutional Animal Welfare Body (OPBA) and authorized by the Italian Ministry of Health (project number approval 480/2021-PR).

### Schwann cells and endoneurial fibroblast cultures

Primary cultures were established from sciatic nerves of 40-day-old WT and CMT1A rats according to published protocols [29]. Briefly, sciatic nerves were dissected under sterile conditions, deprived of perineurium and carefully teased to obtain small fiber bundles which were enzymatically digested for up to 15-18 hours (37°C and 5% CO_2_). Then, samples were mechanically dissociated, centrifuged at 600 g for 5 min at 4°C; the supernatant was removed and the cellular content resuspended in low proliferation medium, consisting of DMEM/F-12 (Gibco #11320033) supplemented with glutamax 2mM (Gibco #35050061), penicillin/streptomycin (Gibco #10378016), and FBS 10% (Gibco #A5256701). Cells were drop-plated in a petri dish coated with poly-L-lysine (Sigma-Aldrich, #P6282) and laminin (Sigma-Aldrich #P2020) overnight. The day after plating, a medium specific for SC expansion (low proliferation medium supplemented with Heregulin 10 ng/mL (Sigma-Aldrich #H7760) and Forskolin 2 µM (Sigma-Aldrich #6886) was gently added. The resulting culture was highly enriched in SCs, but some endoneurial fibroblasts (approximately 10%) were also present; they were isolated and recovered by an immunomagnetic separation using beads coated with the fibroblast specific Thy-1 epitope (Myltenyi Biotec, CD90.1 MicroBeads). Thereafter, SCs and fibroblasts were purified and cultured separately for protein expression studies.

### SYBR Green DNA quantification

Sciatic nerves from P5-P365 WT and CMT1A rats were dissected, carefully weighted and homogenized in 0.25% Triton X-100. Samples were centrifuged (10.000 rcf, 5 min) and the resulting pellets dissolved in TE buffer (Tris HCL 10 mM, EDTA 1 mM, pH 8). Five µL of each sample was loaded into a 96-well plate with 100 µL of SYBR Green solution and 100 µL of TE buffer. After five min at room temperature in the dark, the microtiter plate was read using a fluorescence microplate reader with excitation and emission wavelength at 497 and 520 nm, respectively (Infinite 200 PRO, Tecan Italia Srl). A standard curve was prepared by making serial dilutions (from 10 µg to 2.42 ng) of injected DNA. For each sample, DNA amount was calculated from the difference in fluorescence between the sample and its relative negative control. Resulting values were interpolated with the related standard curve to obtain absolute DNA concentration (µg) [30].

### Nuclear morphometrical analysis

Frozen sections of sciatic nerves from P5-P365 WT and CMT1A rats were counterstained with DAPI and processed as previously described. Nuclei across entire nerve sections were acquired and analyzed using CellProfiler software (v4.2.8) [31]. A set of morphological parameters, including area, compactness, eccentricity, diameter, form factor, mean radius, perimeter, and solidity, was quantified for each nucleus (approximately 19,000 nuclei in total). Graphical representations and ridge plots were generated using Prism and R software.

### Immunofluorescence

To evaluate protein expression, OCT-embedded sciatic nerves were cryosectioned into 10 µm-thick sections, mounted on Superfrost adhesive glass slides (Epredia, J1800AMNZ), and stored at -80 °C until processing. Moreover, immunofluorescences were also conducted on primary SC and endoneurial fibroblasts.

Frozen sections of sciatic nerves and primary cultures were rinsed in Dulbecco’s phosphate-buffered saline (DPBS; Thermo Fisher Scientific, 14190-144), fixed in 4% paraformaldehyde for 20 min, and permeabilized and blocked for 30 min in buffer containing 15% normal goat serum, 0.3% Triton X-100, 20 mM phosphate buffer, and 450 mM NaCl. Sections were then incubated overnight at 4 °C with primary antibodies against MKI67 (Abcam, AB15580, 1:200), JUN (Cell Signaling Technology, 60A8, 1:200), Sox10 (Bio-Techne, AF2864, 1:200), RUNX2 (Cell Signaling Technology, D1L7F, 12556, 1:300), and SMOC2 (Santa Cruz Biotechnology, sc-376104, 1:200).

After washing in DPBS, sections were incubated for 90 min at room temperature with Alexa Fluor 594 or 488 conjugated goat anti-rabbit, goat anti-mouse, and donkey anti-goat IgG secondary antibodies (Thermo Fisher Scientific, A11008, A11032; A11058, used 1:400). Samples were counterstained with DAPI (300 nM), and sections were mounted using Fluoromount aqueous mounting medium.

Whole-section images were acquired using an Olympus PROVIS AX60 microscope equipped with an Olympus DP74 digital camera. Images were acquired at different magnifications using the CellSens image analysis software (Olympus) and analyzed using Fiji [32]. MKI67, JUN, Sox10 and RUNX2^+^ cells were counted and reported as the number of positive cells over the total number of nuclei or as the number of positive cells over the analyzed tissue area.

### Western Blot Analysis

Sciatic nerves from P365 WT and CMT1A rats were homogenized in RIPA lysis buffer (150 mM NaCl, 1% Triton X-100, 0.5% sodium deoxycholate, 0.1% SDS, and 50 mM Tris-HCl, pH 7.5) supplemented with NaF (5 mM), NaVO_4_ (1 mM), phenylmethylsulfonyl fluoride (PMSF, 1 mM), and a protease and phosphatase inhibitor cocktail (Sigma-Aldrich, #P8340; used 1:100).

Protein concentration was determined using a Pierce™ BCA Protein Assay Kit (Thermo Fisher Scientific, #23227) according to the manufacturer’s instructions. Equal amounts of proteins (25 µg per sample) were separated by SDS-PAGE on 4–15% Mini-PROTEAN® TGX Stain-Free™ gels (Bio-Rad, #4568085). Proteins were transferred onto a PVDF membrane by blotting for 1 h at 50 V in Towbin Buffer (25 mM Tris Base, 192 mM Glycine, 20 % (v/v) Methanol, pH 8.5). After transfer, membranes were blocked for 1 h at room temperature in 5% bovine serum albumin (BSA) in Tris-buffered saline containing 0.1% Tween-20 (TBST). Membranes were incubated overnight at 4°C with primary antibodies for the detection of RUNX2 (Cell Signaling Technology, D1L7F, 12556, used 1:1000), or SMOC2 (Santa Cruz Biotechnology, sc-376104, used 1:250) and Vinculin (Sigma-Aldrich, V9264, used 1:10000) used as a loading control. Then, membranes were washed with TBST and incubated for 90 min at room temperature with anti-mouse or anti-rabbit HRP-conjugated secondary antibodies (Santa Cruz Biotechnology, sc-516102 1:5000; sc-2357 used 1:5000). Immunoreactive bands were visualized using Clarity™ Western ECL Substrate (Bio-Rad, #1705060) and imaged with the ChemiDoc™ MP Imaging System. Densitometric analysis was performed using Image Lab software (version 6.1.0, Bio-Rad). The RUNX2 and SMOC2 signal intensities were normalized over the Vinculin for each sample analyzed.

### RNA-seq

Gene expression data relative to WT and CMT1A nerves at 2, 6, and 12 months of age were obtained as described. Briefly, RNA extraction from rat sciatic nerves was obtained by RNAeasy Lipid Tissue Mini Kit (Qiagen). RNA integrity was checked prior to generating RNA-seq libraries by RNA ScreenTape System (Agilent). Sequencing libraries were made using TruSeq Stranded mRNA Library Prep kit (Illumina) following manufacturer’s instructions. For each sample, two technical replicates were generated using 300 ng of total RNA per replicate. Libraries were controlled for correct size using High Sensitivity D1000 ScreenTape System (Agilent). Then, libraries were pooled and quantified by real-time PCR using PhiX Control v3 (Illumina) as a reference. Sequencing was performed using a 75 cycles High Output sequencing kit in a NextSeq platform (Illumina) generating an average of 16 Mreads per technical replicate (32 Mreads per sample). Data were aligned to rn6 Rattus norvegicus genome assembly using STAR v2.7.5c [33]. Transcript quantification was performed using RSEM v1.3.1 [34]. FPKM values generated by RSEM were used to visualize the expression of individual genes using the ggplot2 R package (https://ggplot2.tidyverse.org/) and, after log transformation, for principal component analysis [35]. Raw gene counts were used for differential expression analysis with edgeR [36]. Transcripts with fewer than 0.5 counts per million in all samples were excluded, and library sizes were normalized using the trimmed mean of M-values method [37]. Differential expression between CMT1A and WT nerves was assessed separately at the indicated ages. Immune-cell and cytotoxicity scores were inferred using the MCP-counter method, implemented through the immunedeconv R package [38], [39]. Rat gene symbols were converted to their human orthologues using babelgene R package. Scores were standardized for visualization, and their association with genotype and age was evaluated using linear models including genotype, age, their interaction, and sex as a covariate. Gene-set enrichment analysis was performed using clusterProfiler, with genes ranked according to their log2 fold change in CMT1A versus WT nerves at 12 months [40]. Rat gene sets were retrieved from the Molecular Signatures Database [41]. The injury-associated gene set was obtained from a dataset deposited in the Gene Expression Omnibus (GEO, Accession number GSE177037), containing RNA-seq data from SCs purified from the rat sciatic nerves at different time points after sciatic nerve injury (3, 5, and 7 days). For each time point, differential expression analysis was performed using the R package DESeq2, and for each gene a rank metric was calculated as: log_2_(Fold Change)*-log_10_(p.val) [42]. All genes showing a rank metric higher than 20 at all three time points were included in the gene set (96 genes in total, see **Suppl. Table 1** for the complete list of genes). Positive normalized enrichment scores indicated enrichment in CMT1A nerves. Gene Ontology enrichment analysis of selected differentially expressed gene lists was performed and visualized using the ClueGO Cytoscape plugin [43].

### Statistical analysis and data visualization

For each experiment, the statistical test used is reported in the figure legends. The threshold for statistical significance was set at p < 0.05. Unless otherwise stated, all data are represented as mean ± SEM. In the figure legends, mean ± SD is also reported alongside the exact p-value for each comparison. GraphPad Prism software version 10 (GraphPad Software Inc., San Diego, CA, USA) was used to perform the statistical comparisons and to generate the plots. All figures were prepared using Adobe Illustrator 2023 (Adobe, San Jose, CA, USA).

## Results

*CMT1A nerves exhibit persistent cellular proliferation in adult rats*.

Hypercellularity in experimental and human CMT1A, especially during development, has been widely reported in the literature; however, its persistence and progression over time remain incompletely characterized. We combined SYBR Green DNA quantification with histological analysis of nuclei and we found that CMT1A sciatic nerves contained a higher amount of DNA and maintained a higher nuclear density compared with WT controls, even though nuclear density progressively decreased over time in both genotypes, likely reflecting nerve growth and tissue expansion (**Fig. 1A-C**). In particular, CMT1A sciatic nerves showed higher DNA amount and nuclei number compared to WT littermates starting at P20 and reaching a threefold gain at P365 **(Fig. 1A-C**). Furthermore, the distribution analysis of nuclear size in WT and CMT1A sciatic nerves revealed the persistence of larger nuclei in CMT1A nerves, a feature commonly associated with active proliferation across multiple cell types (**Fig. 1D**) [44]. Notably, this hypercellularity phenotype is not restricted to the sciatic nerve, but likely extends to the entire peripheral nervous system, as we also observed increased cell number in both the femoral and saphenous nerves, representing predominantly motor and sensory nerves, respectively (**Suppl. Fig. 1**).

**Figure 1.**
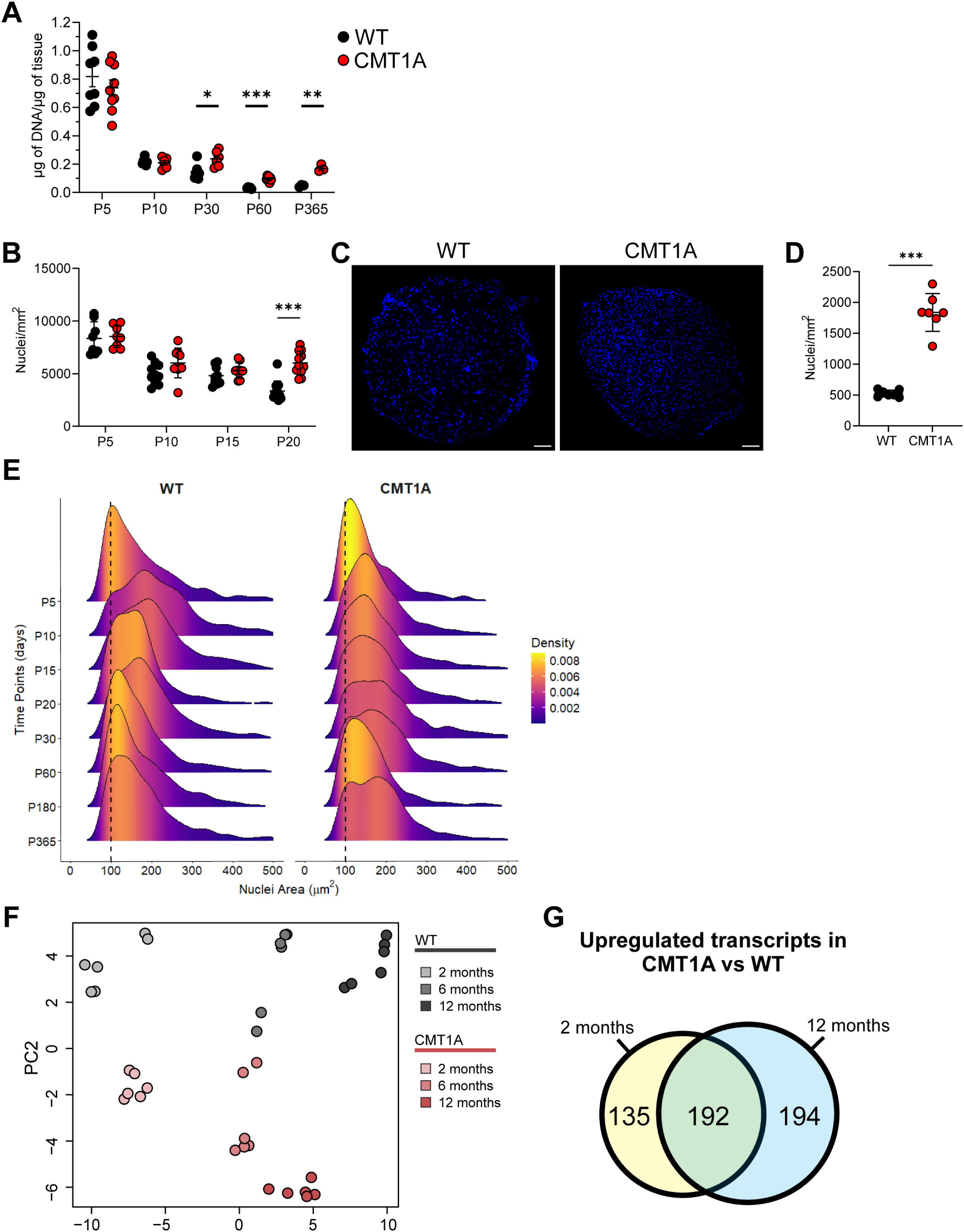
CMT1A nerves exhibit persistent cellular proliferation in adult rats. (A) DNA content of WT and CMT1A rat sciatic nerves at P5, P10, P30, P60, and P365, normalized over μg of tissue. (Mean ± SD - WT vs CMT1A: P5: 0.82 ± 0.2 vs 0.74 ± 0.17, p = ns, N = 8 mice per genotype; P10: 0.22 ± 0.02 vs 0.21 ± 0.04, p = ns, N = 7 mice per genotype; P30: 0.14 ± 0.05 vs 0.24 ± 0.05, p < 0.05, N = 7 mice per genotype; P60: 0.03 ± 0.005 vs 0.10 ± 0.02, p < 0.001, N = 7 mice per genotype; P365: 0.05 ± 0.008 vs 0.17 ± 0.02, p < 0.01, N = 3 mice per genotype). p-values were calculated using multiple two-tailed unpaired t tests corrected for multiple comparisons using the Holm-Šídák method. (B) Nuclei density in sciatic nerves sections from WT and CMT1A rats at P5, P10, P15, and P20. (Mean ± SD - WT vs CMT1A, N = 8 mice per genotype and per time point: P5: 8349 ± 1575 vs 8525 ± 1041, p = ns; P10: 4990 ± 999 vs 6013 ± 1405, p = ns; P15: 4812 ± 891 vs 5297 ± 739, p = ns; P20: 3350 ± 967 vs 6028 ± 1116, p < 0.001.) For each mouse, values were averaged across at least 3 sections. p-values were calculated using multiple two-tailed unpaired t tests corrected for multiple comparisons using the Holm-Šídák method. (C) Sciatic nerves sections from WT and CMT1A adult (P365) rats, stained with DAPI. Scale bars: 100 μm. (D) Nuclei density in sciatic nerves sections from adult (P365) WT and CMT1A rats. (Mean ± SD - WT vs CMT1A, N = 8 mice per genotype: 525 ± 56.1 vs 1839 ± 307, p < 0.001). For each mouse, values were averaged across at least 3 sections. p-values were calculated using two-tailed unpaired t test with Welch’s correction. (E) Ridgeplot showing the frequency density of differently sized nuclei in WT and CMT1A sciatic nerves at P5, P10, P15, P20, P30, P60, P180, and P365. (F) Principal component analysis of bulk RNA-seq profiles from WT and CMT1A sciatic nerves collected at 2, 6, and 12 months of age, restricted to genes differentially expressed between 2- and 12-month-old WT nerves. (G) Venn diagram showing transcripts significantly upregulated in CMT1A compared with WT nerves at 2 and 12 months. A total of 192 transcripts were shared between time points, whereas 135 and 194 were specific to 2 and 12 months, respectively. Data are presented as mean ± SEM. * = p < 0.05, ** = p < 0.01, *** = p < 0.001

To gain molecular insight into this persistent tissue event, we performed a bulk RNA sequencing analysis on sciatic nerves collected from 2-, 6-, and 12-month-old rats. Unsupervised principal component analysis (PCA) showed a clear segregation between WT and CMT1A genotypes at all analyzed time points, whereas age-related differences were comparatively less pronounced (**Suppl. Fig. 2A**). A subsequent PCA restricted to genes associated with physiological aging (i.e. differentially expressed between 2- and 12-month-old nerves) revealed a genotype-dependent divergence that exacerbates with age (**Fig. 1E**). In line with this progressive divergence, differential expression analysis identified a larger number of differentially expressed genes (DEGs) between CMT1A and WT nerves at 12 months than at 2 months of age. Overall, 192 DEGs were shared between the two time points, whereas 135 and 194 were specific to the early and late stages, respectively (**Fig. 1F**). These findings suggest that a core disease-associated transcriptional signature is maintained over time, while additional alterations emerge with age.

Consistent with our previous findings, these nerves displayed significant enrichment of gene signatures associated with cell proliferation, protein biosynthesis, and MYC-regulated pathways (**Fig. 2A**). Among the most highly expressed transcripts were DNA topoisomerase II alpha (Top2a), Aurora kinase B (*Aurkb*), centromere protein A (*Cenpa*), and the proliferation marker Ki-67 (*Mki67*), collectively supporting sustained activation of proliferative programs in CMT1A pathology (**Fig. 2B**). Actually, these genes followed a similar temporal pattern in both genotypes: although their expression decreased with age, their levels remained consistently higher in CMT1A compared with WT at all time points.

**Figure 2.**
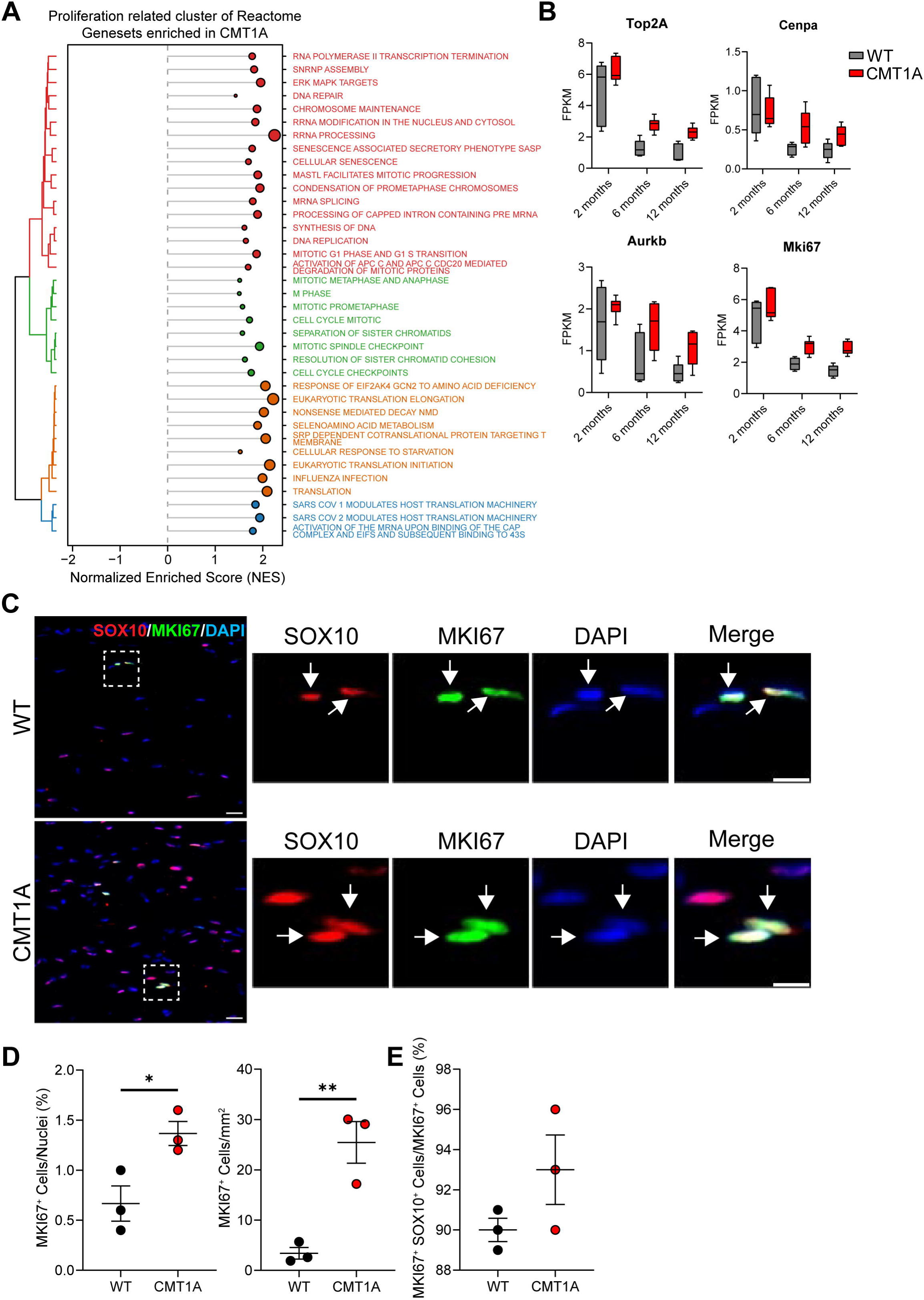
Sequencing and Histological Data confirm persistent cellular proliferation in CMT1A nerves. (A) Gene-set enrichment analysis of cell-type signatures in 12-month-old CMT1A versus WT nerves. Positive normalized enrichment scores (NES) indicate signatures enriched in CMT1A, whereas negative NES values indicate signatures enriched in WT. Colors denote clusters of related signatures. (B) Expression levels in WT and CMT1A nerves of 4 genes known to be key regulators of cell proliferation. (C) Sciatic nerve sections from 12-month-old WT and CMT1A rats immunostained for SOX10 and MKI67, and counterstained with DAPI. Panels on the right represent zoomed images of the region highlighted by the dashed square in the panel on the left. Scale bars: 20 μm in the images on the left, 10 μm in the zoomed images. (D) Density of MKI67^+^ cells in sciatic nerve sections from 12-month-old WT and CMT1A rats, shown as the percentage of MKI67^+^ cells over total nuclei (left), and as number of MKI67^+^ cells per mm^2^ (right). (Mean ± SD - WT vs CMT1A, N = 3 mice per genotype: MKI67^+^cells/nuclei: 0.67 ± 0.14 vs 1.24 ± 0.20, p < 0.05; MKI67^+^ cells/mm^2^: 3.4 ± 2.02 vs 22.3 ± 6.13, p < 0.05). For each mouse, values were averaged across at least 3 sections. p-values were calculated using two-tailed unpaired t test with Welch’s correction. (E) Percentage of SOX10^+^ MKI67^+^ cells over the total number of MKI67^+^ cells. (Mean ± SD - WT vs CMT1A, N = 3 mice per genotype: 90 ± 1 vs 93 ± 3, p = ns). For each mouse, values were averaged across at least 3 sections. p-values were calculated using two-tailed unpaired t test with Welch’s correction. Data are presented as mean ± SEM.* = p < 0.05, ** = p < 0.01.

In agreement with transcriptional data, protein analysis revealed an approximately threefold increase in MKI67^+^ nuclei in 12-month-old CMT1A sciatic nerves relative to WT controls, indicating that a consistent fraction of the observed hypercellularity reflects ongoing cell cycle activity (**Fig. 2C,D**). Moreover, the vast majority of MKI67^+^ cells (∼90%) also expressed SOX10, the key transcription factor governing SC differentiation, indicating that proliferating cells predominantly belong to the SC lineage (**Fig. 2E**).

### Cell-Type Signature Enrichment Reveals Immature and Mesenchymal-Like Transcriptional Programs in CMT1A Nerves

To further dissect the transcriptional programs associated with aberrant proliferation in CMT1A, we used a deconvolution approach to estimate immune cell abundance and cytotoxic activity from sciatic nerve transcriptomes. None of the inferred immune-related scores showed a significant genotype-by-age interaction, suggesting that CMT1A nerve pathology is not associated with a progressive immune cell infiltration or accumulation (**Fig. 3A**).

**Figure 3.**
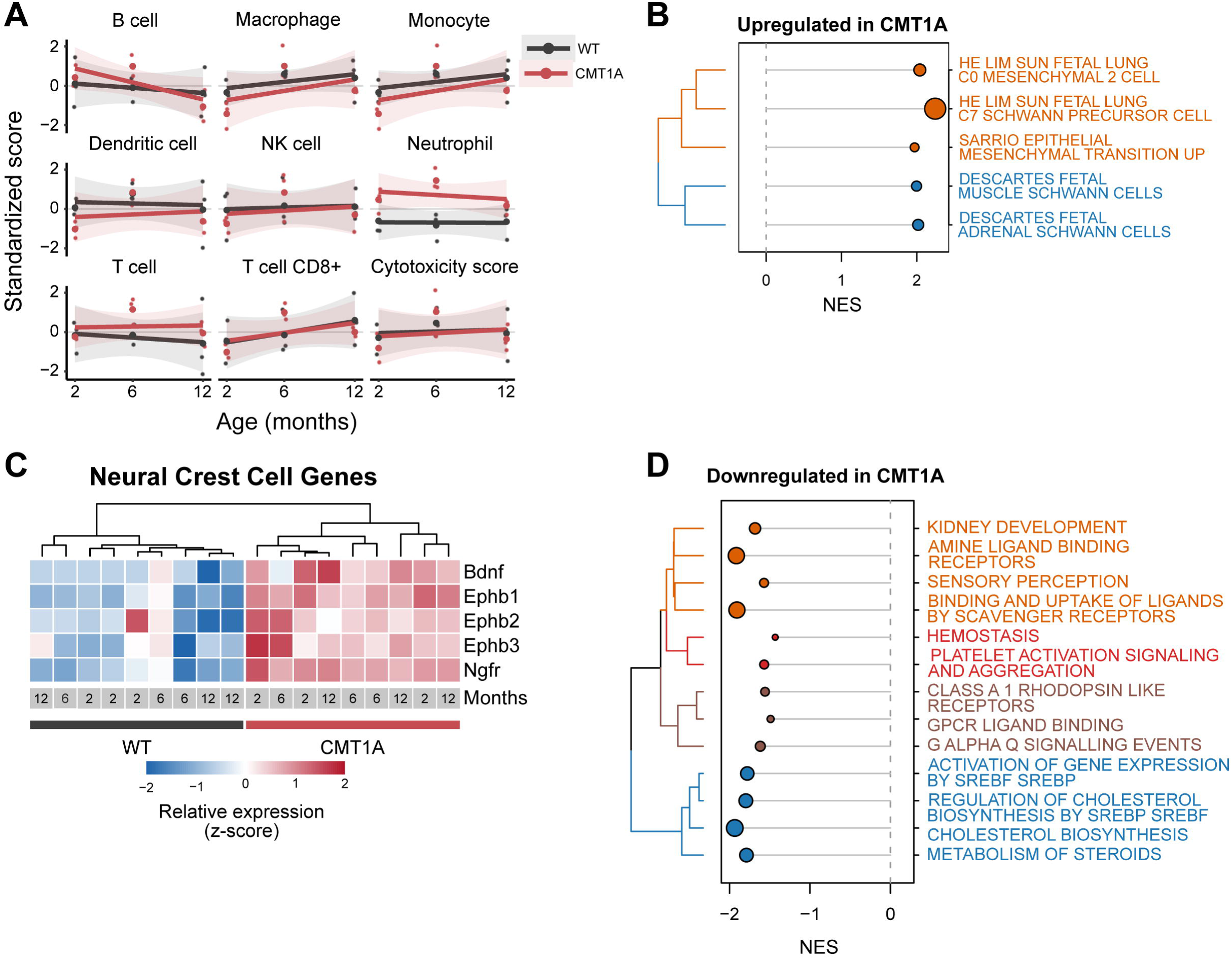
Sequencing data show upregulation of mesenchymal gene signatures in CMT1A nerves. (A) Immune-cell and cytotoxicity scores inferred from bulk RNA-seq data and shown as standardized scores across age and genotype. Points represent individual samples; lines and shaded areas show the fitted trend and 95% confidence interval. (B) Gene-set enrichment analysis of Reactome proliferation-related pathways enriched in CMT1A nerves. Dot size reflects gene-set size and colors denote clusters of related pathways. (C) Heatmap of leading-edge genes from the neural crest cell genes set. Expression values are row-scaled z-scores; samples are grouped by genotype and annotated by age. (D) Gene-set enrichment analysis of pathways negatively enriched in CMT1A nerves; colors denote clusters of related pathways.

Cell-type signature enrichment analysis revealed that CMT1A nerves were enriched in transcriptional profiles associated with fetal SCs and SC precursors, together with fetal mesenchymal and epithelial-to-mesenchymal transition signatures (**Fig. 3B**). Collectively, these profiles point toward the activation of immature and mesenchymal-like transcriptional programs in CMT1A nerves.

In support of this interpretation, we identified a subset of transcripts upregulated in CMT1A nerves that is associated with neural crest cells, the developmental progenitors of SCs, which retain the capacity to differentiate into both neural and mesenchymal lineages (**Fig. 3C**) [45], [46].

Conversely, CMT1A nerves showed a downregulation of pathways involved in lipid, cholesterol, and steroid biosynthesis, consistent with the metabolic programs required for mature SC function and myelin maintenance (**Fig. 3D**).

To characterize the hypercellularity observed in CMT1A, we focused specifically on the top 70 genes significantly upregulated in 12-month-old CMT1A rats compared with WT controls (**Suppl. Table 2**). Gene Ontology analysis on this set of genes revealed that the most enriched biological categories included calcium ion sequestration, ERBB signalling pathways, and embryonic cranial skeleton morphogenesis (**Suppl. Fig. 2B** and **Suppl. Table 3**). Given the prominence of skeletal morphogenesis related genes, we further explored transcripts linked to mesenchymal differentiation, consistent with the transition phenotype emerging from our analyses.

Among the upregulated genes were *Retinol dehydrogenase 10* (*Rdh10*), *Methylenetetrahydrofolate dehydrogenase 1-like* (*Mthfd1l*), and *RUNX family transcription factor 2* (*RUNX2*) (**Fig. 4A**). RUNX2 was of particular interest due to its established role in stem cell regulation and its emerging involvement in SC responses to sciatic nerve injury, including pathways mediated by JUN [27], [47].

**Figure 4.**
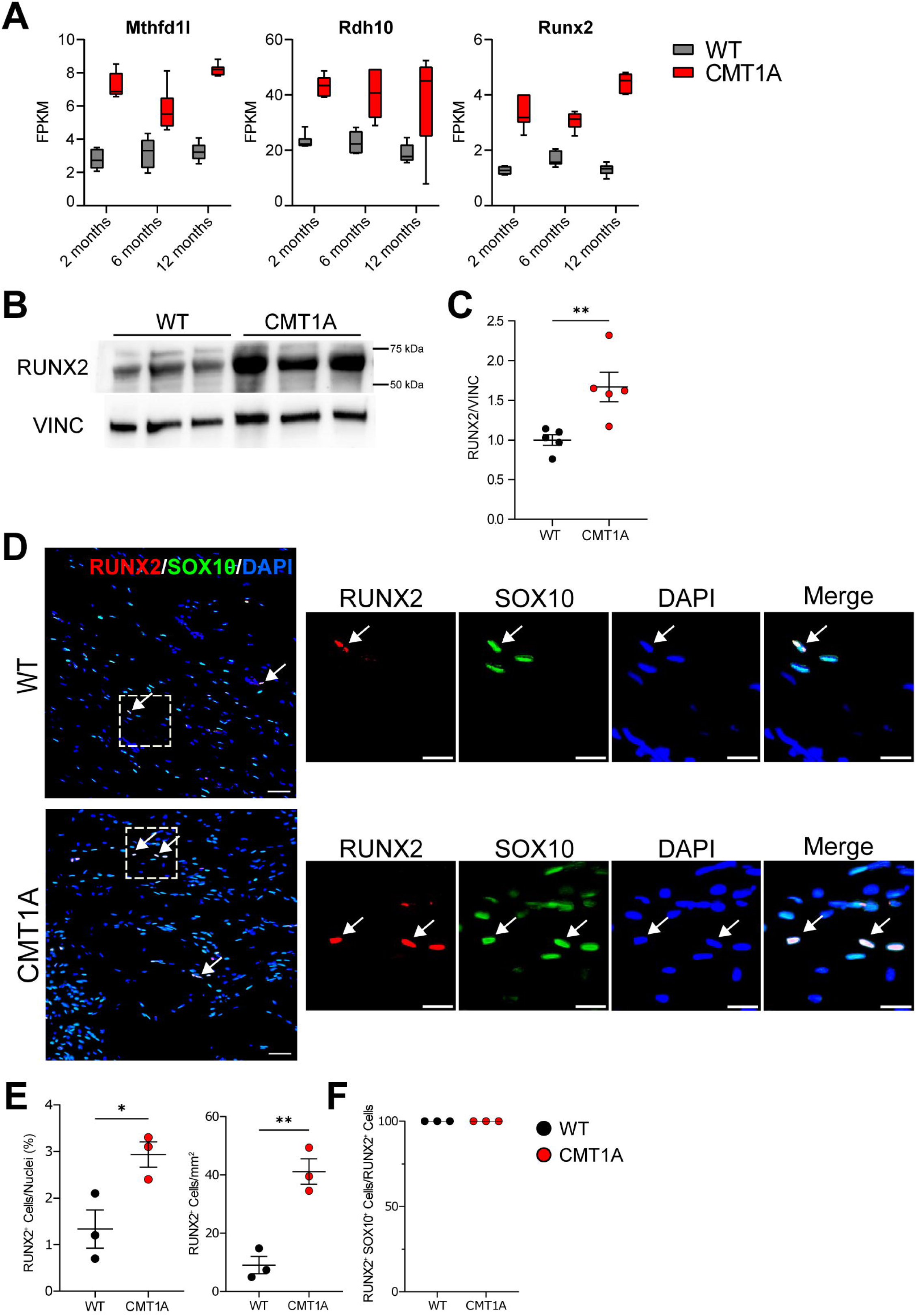
RUNX2 is upregulated in CMT1A sciatic nerves. (A) Expression levels in WT and CMT1A nerves of 3 genes known to be key regulators of mesenchymal-like progenitors (B-C) Western blot (B) and densitometric analysis (C) of RUNX2 expression in sciatic nerves from 12-month-old WT and CMT1A rats. Vinculin was used as a loading control. (Mean ± SD - WT vs CMT1A, N = 5 mice per genotype: 1 ± 0.15 vs 1.67 ± 0.41, p < 0.05). p-values were calculated using two-tailed unpaired Student’s t test. (D) Sciatic nerve sections from 12-month-old WT and CMT1A rats immunostained for SOX10 and RUNX2, and counterstained with DAPI. Panels on the right represent zoomed images of the region highlighted by the dashed square in the panel on the left. Scale bars: 20 μm. (E) Density of RUNX2^+^ cells in sciatic nerve sections from 12-month-old WT and CMT1A rats, shown as the percentage of RUNX2^+^ cells over total nuclei (left), and as number of RUNX2+ cells per mm^2^ (right). (Mean ± SD - WT vs CMT1A, N = 3 mice per genotype: RUNX2^+^ cells/nuclei: 1.33 ± 0.71 vs 2.93 ± 0.47, p < 0.05; RUNX2^+^ cells/mm^2^: 9.05 ± 5.1 vs 41.13 ± 7.54, p < 0.01). For each mouse, values were averaged across at least 3 sections. p-values were calculated using two-tailed unpaired t test with Welch’s correction. (F) Percentage of SOX10+ RUNX2+ cells over the total number of RUNX2+ cells. (Mean ± SD - WT vs CMT1A, N = 3 mice per genotype: 100 ± 0 vs 100 ± 0). For each mouse, values were averaged across at least 3 sections. p-values were calculated using two-tailed unpaired t test with Welch’s correction. Data are presented as mean ± SEM. * = p < 0.05, ** = p < 0.01.

### Emergence of RUNX2-Positive Schwann Cells and Altered JUN Associated Injury Responses in CMT1A Nerves

We confirmed RUNX2 overexpression in CMT1A nerves using Western Blot (**Fig. 4B,C**). Moreover, immunohistological analyses revealed that pathological nerves displayed a marked increase in the number of RUNX2^+^ nuclei compared with controls (**Fig. 4D,E**). Then, we assessed the colocalization of RUNX2 with SOX10. We observed complete overlap between RUNX2^+^ and SOX10^+^ cells, indicating that RUNX2 expression is confined to cells belonging to the SC lineage in both WT and CMT1A sciatic nerves (**Fig. 4F**).

Given the established role of RUNX2 in nerve injury responses, we next investigated the expression of JUN, the master regulator of SC injury reprogramming, and SOX10. Notably, transcripts of both genes were significantly increased in CMT1A sciatic nerves (**Fig. 5A**). These findings were confirmed at the protein level by immunofluorescence analysis. (**Fig. 5B-F**).

**Figure 5.**
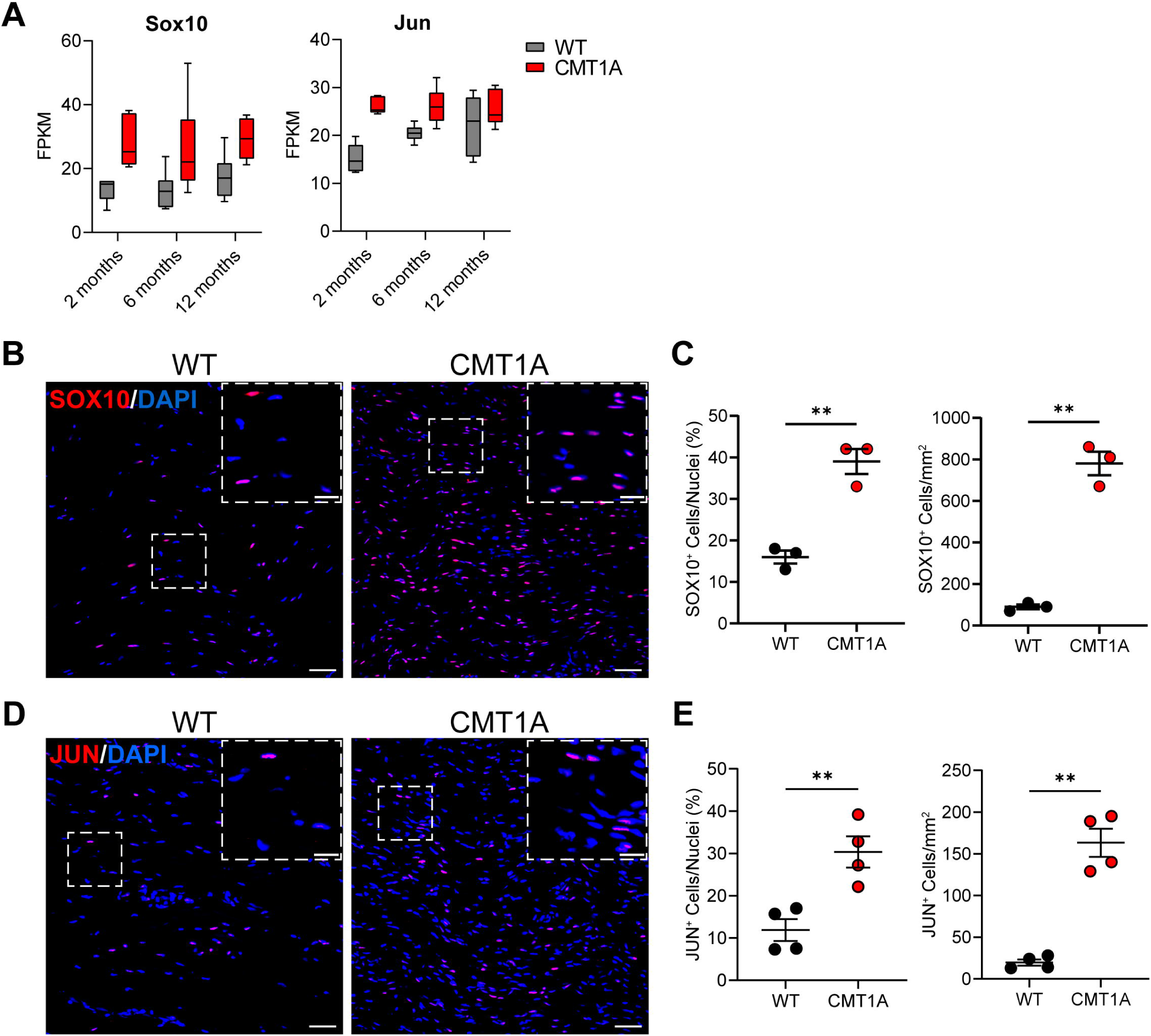
SOX10^+^ and JUN^+^ cell number is increased in CMT1A sciatic nerves. (A) Expression levels in WT and CMT1A nerves of the Schwann cell (SC) lineage marker *Sox10* (left), and of the injury SC marker *Jun* (right). (B) Sciatic nerve sections from 12-month-old WT and CMT1A rats immunostained for SOX10, and counterstained with DAPI. Inserts on the top right corner of each image represent zoomed images of the region highlighted by the dashed square. Scale bars: 20 μm (C) Density of SOX10^+^ cells in sciatic nerve sections from 12-month-old WT and CMT1A rats, shown as the percentage of SOX10^+^ cells over total nuclei (left), and as number of SOX10^+^ cells per mm^2^ (right). Mean ± SD - WT vs CMT1A, N = 3 mice per genotype: SOX10^+^ cells/nuclei: 16 ± 3 vs 39 ± 5, p < 0.01; SOX10^+^ cells/mm^2^: 0.09 ± 0.02 vs 0.78 ± 0.10, p < 0.01). For each mouse, values were averaged across at least 3 sections. (D) Sciatic nerve sections from 12-month-old WT and CMT1A rats immunostained for JUN, and counterstained with DAPI. Inserts on the top right corner of each image represent zoomed images of the region highlighted by the dashed square. Scale bars: 20 μm. (E) Density of JUN^+^ cells in sciatic nerve sections from 12-month-old WT and CMT1A rats, shown as the percentage of JUN^+^ cells over total nuclei (left), and as number of JUN^+^ cells per mm^2^ (right). For each mouse, values were averaged across at least 3 sections. p-values were calculated using two-tailed unpaired t test with Welch’s correction. Mean ± SD - WT vs CMT1A, N = 3 mice per genotype: JUN^+^cells/nuclei: 11 ± 5 vs 30 ± 7, p < 0.01; JUN^+^ cells/mm^2^: 20 ± 7 vs 163 ± 34, p < 0.01). p-values were calculated using two-tailed unpaired t test with Welch’s correction. Data are presented as mean ± SEM. ** = p < 0.01.

To further dissect this response, we performed colocalization analysis of JUN and SOX10. SOX10^+^/JUN^+^ cells were detected almost exclusively in CMT1A sciatic nerves (yellow fraction of the bars). In addition, SOX10^+^/JUN^-^(red) and SOX10^-^/JUN^-^populations (blue) were also increased in CMT1A (**Fig. 6A,B**). The enrichment of SOX10^+^/JUN^+^ cells is consistent with the emergence of injury-associated SCs, commonly referred to as Büngner cells [48].

**Figure 6.**
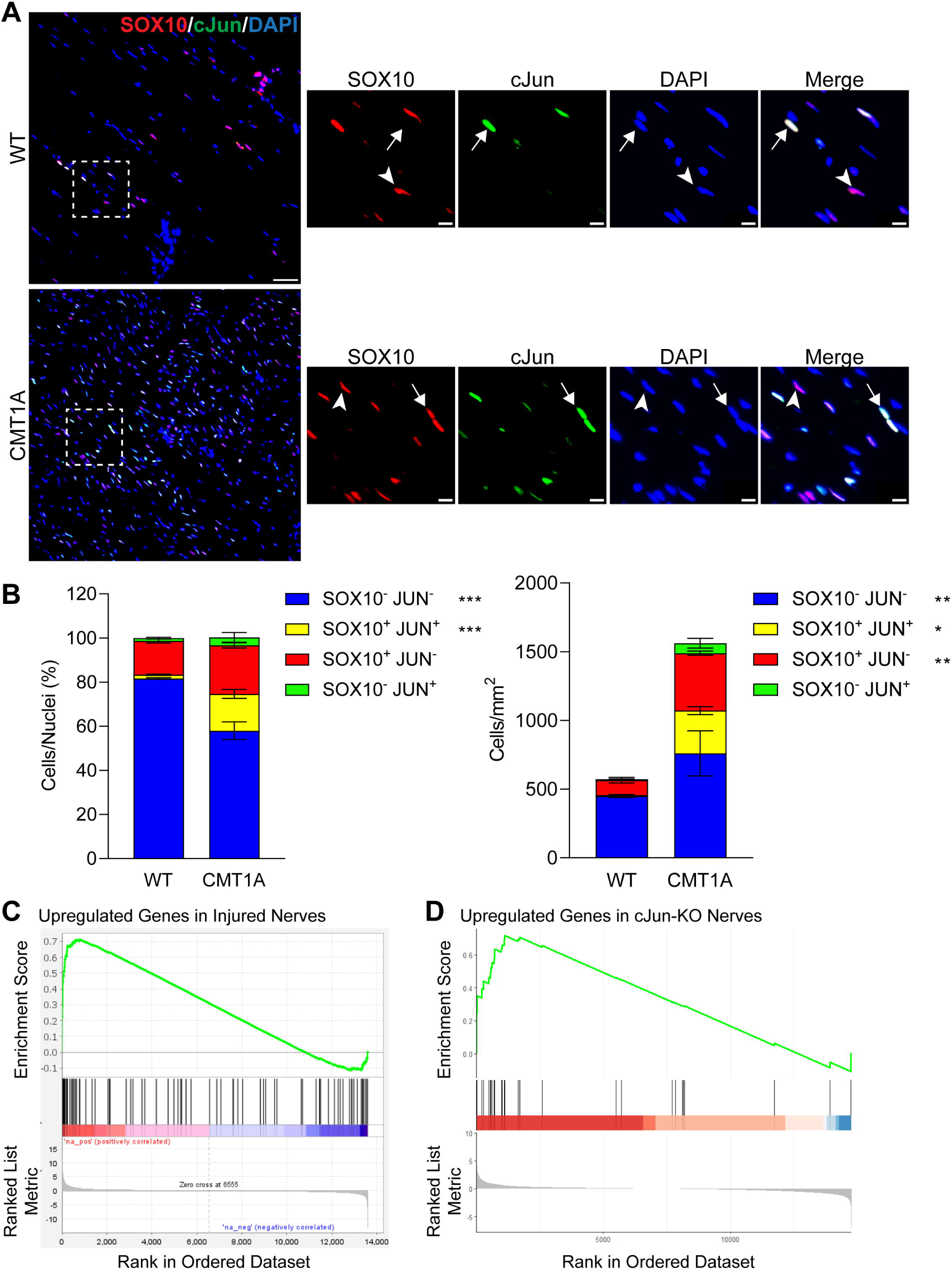
An injury response-like transcriptional program is defectively activated in CMT1A sciatic nerves. (A) Sciatic nerve sections from 12-month-old WT and CMT1A rats immunostained for SOX10 and JUN, and counterstained with DAPI. Panels on the right represent zoomed images of the region highlighted by the dashed square in the panel on the left. Scale bars: 20 μm in the images on the left, 10 μm in the zoomed images. (B) Density of SOX10^-^JUN^-^, SOX10^+^;JUN^+^, SOX10^+^ JUN^-^, and SOX10^-^JUN^+^ cells, reported as percentage over total nuclei (left), and as total number of cells per mm^2^ (right). (Mean ± SD - WT vs CMT1A, N = 3 mice per genotype: SOX10^-^JUN^-^cells/nuclei: 82 ± 1 vs 58 ± 7, p < 0.001; SOX10^+^ JUN^+^ cells/nuclei: 2 ± 1 vs 17 ± 4, p < 0.001; SOX10^+^ JUN^-^cells/nuclei: 16 ± 2 vs 22 ± 2, p = ns; SOX10^-^JUN^+^ cells/nuclei: 1 ± 1 vs 4 ± 4, p = ns; SOX10^-^JUN^-^cells/mm^2^: 450 ± 19 vs 760 ± 283, p < 0.001; SOX10^+^ JUN^+^ cells/mm^2^: 7 ± 5.3 vs 312 ± 50, p < 0.05; SOX10^+^ JUN^-^cells/mm^2^: 108 ± 35 vs 417 ± 25, p < 0.001; SOX10^-^JUN^+^ cells/mm^2^: 8 ± 3 vs 73 ± 61, p = ns). For each mouse, values were averaged across at least 3 sections. p-values were calculated using two-way ANOVA followed by a post-hoc Šidák multiple comparisons test. (C, D) Gene-set enrichment analysis performed on the CMT1A versus WT differential-expression profile using gene sets comprising genes upregulated following peripheral nerve injury (C) and genes upregulated in JUN-deficient injured nerves (D). Genes were ranked according to differential expression between CMT1A and WT nerves at 12 months; black vertical bars indicate the positions of genes belonging to each gene set, and the green line represents the running enrichment score. Positive values for enrichment score correspond to enrichment in CMT1A. Data are presented as mean ± SEM. * = p < 0.05, ** = p < 0.01, *** = p < 0.001.

Gene Set Enrichment Analysis (GSEA) further revealed a significant upregulation of genes associated with the injury response in CMT1A (**Fig. 6 C**) [49]. To further characterize this injury-prone population, we interrogated our RNA-seq dataset using a set of genes previously identified as dysregulated in JUN knockout mice [47]. Surprisingly, genes reported to be upregulated in JUN–deficient nerves, where the injury response was defeated, were enriched in CMT1A (**Fig. 6** D).

Together, these findings support the presence of an activated yet transcriptionally dysregulated repair program in CMT1A, suggesting a chronic injury-like response that is initiated but functionally impaired.

Finally, given the increase of SOX10^-^cell population in CMT1A nerves, we examined our RNA-seq dataset for established markers of endoneurial fibroblasts, a major non-SC type in peripheral nerve tissue (**Suppl. Fig. 3C**) [50]. Only a limited number of fibroblast specific genes were differentially expressed in CMT1A nerves, possibly reflecting the fact that many canonical markers are associated with fully differentiated fibroblasts, a state that may not completely emerge under CMT1A condition. Among the identified candidates, we focused on SMOC2, an extracellular matrix (ECM) protein. In our dataset, SMOC2 was markedly upregulated in CMT1A sciatic nerves at both the mRNA and protein levels (**Fig. 7A-C**). Immunofluorescence analysis further demonstrated that SMOC2 expression was restricted to endoneurial fibroblasts and absent in SCs (**Fig. 7D**). These findings point to the activation of a fibroblast-specific transcriptional program that has not previously been described in CMT1A. This observation supports the hypothesis that aberrant endoneurial fibroblast function contributes to the pathological tissue remodeling observed in affected nerves.

**Figure 7.**
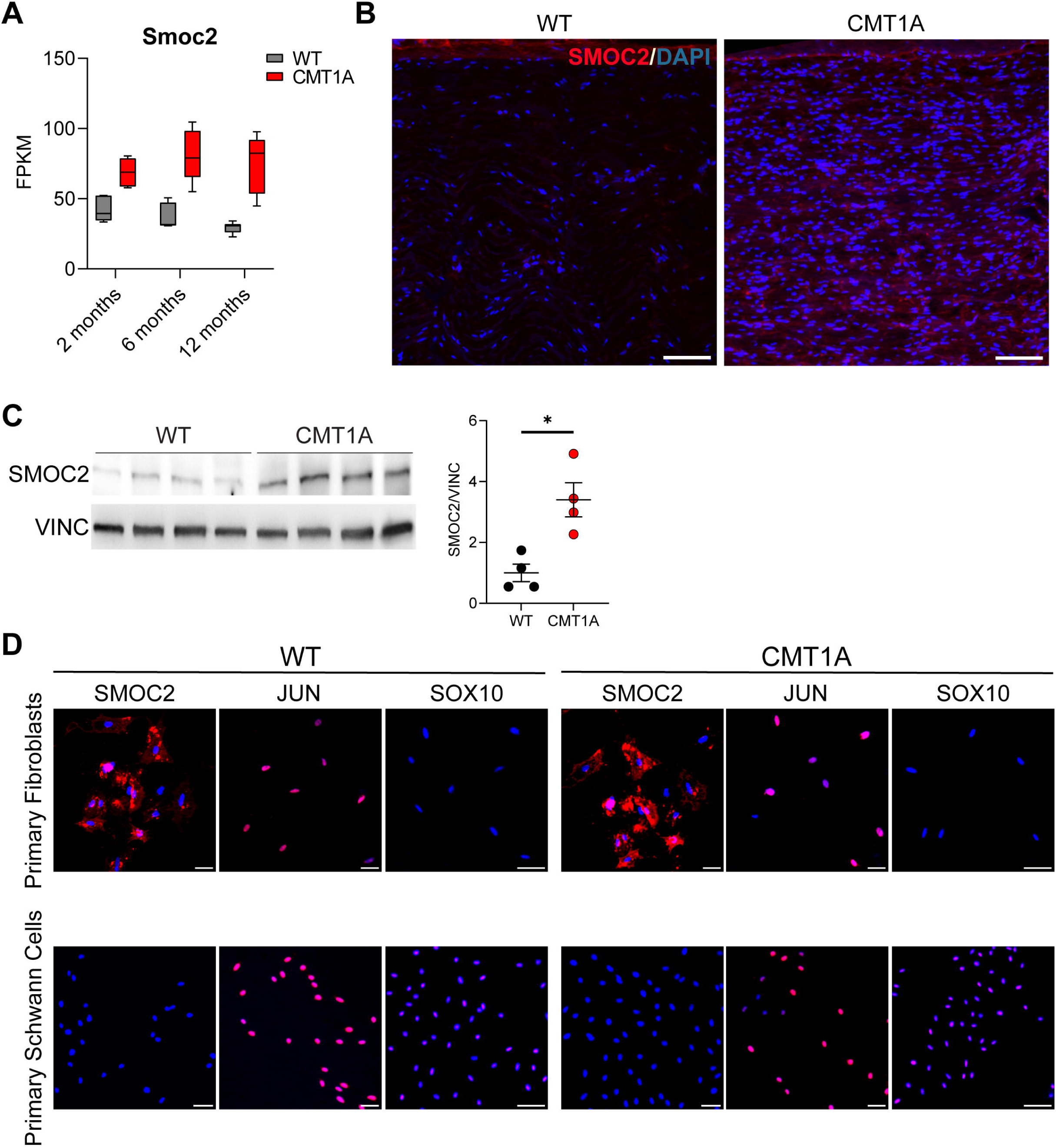
The fibroblast-specific SMOC2 protein is upregulated in CMT1A nerves. (A) Expression levels in WT and CMT1A nerves of *Smoc2*. (B) Sciatic nerve sections from 12-month-old WT and CMT1A rats immunostained for SMOC2, and counterstained with DAPI. Scale bars: 50 μM. (C) Western blot (left) and corresponding densitometric analysis (right) of SMOC2 expression in sciatic nerves from 12-month-old WT and CMT1A rat. Vinculin was used as a loading control. (Mean ± SD - WT vs CMT1A, N = 4 mice per genotype: 1 ± 0.58 vs: 3.2 ± 1.12, p < 0.05). p-values were calculated using two-tailed unpaired t test with Welch’s correction. (D) Purified fibroblast and SC cultures from WT and CMT1A rats, immunostained for SMOC2, JUN, or SOX10, and counterstained with DAPI. Scale bars: 20 μM. Data are shown as mean ± SEM. * = p < 0.05.

## Discussion

### Chronic Proliferative State Underlies CMT1A Neuropathy

In this study, we demonstrate the presence of remarkable and sustained proliferative activity in the sciatic nerves of CMT1A rats throughout their lifespan. To our knowledge, this is the first report showing that increased proliferation and hypercellularity in CMT1A nerves persists up to 12 months of age, well beyond the canonical developmental period. Notably, proliferative activity is initiated before postnatal day 20 in both WT and CMT1A rats. However, while physiologically declining over time, it remains abnormally high in adult CMT1A nerves. This pattern is further supported by the expression profiles of proliferation-associated genes, including *Mki67*. The limited portion of proliferating cells observed (∼3%), despite the markedly increased cell density, may reflect features commonly associated with tissue stem or progenitor cell populations. Such cells typically exhibit a slow-cycling proliferative behavior, which enables controlled lineage expansion while preserving genomic stability and long-term multipotency [45], [51]. Interestingly, proliferation levels at 2 months of age remain relatively high even in WT animals, suggesting that peripheral nerve growth and maturation extend longer than previously appreciated [11]. These findings imply that the therapeutic window for CMT1A might broaden than traditionally assumed.

A critical issue concerns the cellular source underlying the persistent proliferative activity observed in CMT1A. Our RNA-seq data are not consistent with macrophage or immune infiltration. Given the threefold increase in nuclei number, also onion bulb formation does not seem a plausible primary contributor. Indeed, analysis of cross-sectional sciatic nerve sections revealed only approximately 10–15 such onion bulbs per section [21]. Furthermore, onion bulb–associated supernumerary cells are negative for cyclin D1, a marker of active proliferation that is upregulated in CMT1A and was also increased in our RNA-seq dataset [15]. Cyclin D1 expression was predominantly localized to myelinating SCs along axons. Additionally, in several CMT1A rodent models, onion bulbs are either absent or appear only after several months of age, well after the onset of hypercellularity [14], [19], [20]. Collectively, these observations indicate that onion bulb–associated cells contribute only minimally to the overall increase in cell number and density.

Regarding inflammatory involvement, our data do not support a substantial contribution of classical inflammatory signatures. Moreover, we demonstrate that actively dividing cells belong almost exclusively to the Schwann cell lineage, thereby excluding a substantial contribution of other cell types to the proliferation observed in CMT1A nerves. Previous studies have reported a modest increase in resident macrophages in CMT1A nerves; however, these latter are limited to few cells per area and thereby insufficient to account for the extensive hypercellularity observed throughout life [23], [52].

Another relevant aspect concerns the role of PMP22 in regulating cell proliferation. PMP22 has been described as a growth arrest–specific factor in both *in vitro* and *in vivo* contexts, although other studies suggest a role in promoting proliferation, including oncogenic functions in certain human cancers [26], [53], [54]. Notably, the human *PMP22* gene produces six alternatively spliced transcript variants, driven by two distinct promoters whose usage is altered in CMT1A, supporting potentially distinct roles in myelination and cell cycle regulation [55], [56]. To date, there is no direct evidence linking PMP22 dysregulation to the proliferative phenotype observed in CMT1A.

Importantly, increased proliferation and hypercellularity are not unique to CMT1A but have also been described in other hereditary and toxic neuropathies [57], [58], [59]. This suggests that dysregulation of the SC cycle may represent a common pathogenic mechanism and a potential target for therapeutic intervention.

### RUNX2-Linked Schwann Cell Injury Programs and Fibroblast-Driven Remodeling in CMT1A

A second major finding of this study is the demonstration of increased expression of the transcription factor RUNX2 concomitant with sustained proliferative activity in CMT1A. RUNX2, together with RDH10 and MTHFDL1, ranks among the most upregulated genes identified in our RNA-seq dataset and is associated with mesenchymal and neural crest transcriptional signatures. These results are consistent with a delay in tissue maturation and with the presence of a proliferative cell population within CMT1A sciatic nerves [11], [15].

Notably, RUNX2, widely recognized as a master regulator of osteogenesis, has also been implicated in the response to sciatic nerve injury [47], [60]. In this context, previous work has linked the emergence of an immature/mesenchymal phenotype to injury-induced cellular programs, demonstrating that acquisition of a fully reparative phenotype arises from a complex dedifferentiation process that includes an epithelial–mesenchymal transition (EMT) phase [61], [62].

Furthermore, it is reported that immature SCs do not exhibit a strictly defined phenotype but rather comprise a heterogeneous population biased toward multiple differentiation trajectories [63]. This heterogeneity may account for the lack of consistent modulation of specific lineage markers observed in our RNA-seq data, as well as in previously published datasets [50], [61], [64], [65].

The role of RUNX2 in the nerve injury response is increasingly recognized [66]. Following sciatic nerve injury, RUNX2 expression has been reported to be self-reinforcing and regulated by JUN activity, particularly during the early phases of the injury response. In this context, RUNX2 has been proposed to exert structural and metabolic effects, including the suppression of lipid metabolism, which is thought to contribute to Schwann cell reprogramming after injury [67]. Adding complexity, Büngner repair cells exhibit at least three distinct transcriptional profiles: proliferating, pro-myelinating, and transitional, all of which express JUN. Notably, RUNX2 is predominantly enriched in pro-myelinating repair cells, suggesting a partially distinct role compared to JUN in the injury response and, consequently, in CMT1A-associated hypercellularity [27], [67]. Moreover, recent evidence indicates that transcription factors such as TFEB/3, which are involved in the SC injury response and may act upstream of RUNX2, do not influence JUN levels [68].

Our data are consistent with, and extend, this framework, suggesting that RUNX2 may exert a more restricted and partially independent role rather than functioning exclusively as a downstream effector of JUN. Although RUNX2 and JUN expression were not directly assessed within the same cells, we observed that RUNX2^+^ nuclei were markedly less abundant than JUN^+^ nuclei in CMT1A nerves. Based on this difference in prevalence, it can be inferred that RUNX2-expressing cells likely represent only a minor fraction of the broader JUN^+^ population. This observation raises the possibility that RUNX2 marks a distinct, limited subset of injury-responsive SCs, potentially differing in temporal or spatial dynamics from classical, fully developed Büngner cells. Such a population may contribute to the altered injury-like response observed in CMT1A neuropathy, although further studies will be required to directly test this hypothesis.

The precise mechanisms by which RUNX2 contributes to the structural organization of CMT1A nerves remain unclear. In contrast, it is well established that JUN knockout in CMT1A exacerbates disease severity, and that aging is associated with reduced JUN levels and a diminished injury response capacity, highlighting its protective role [69], [70]. Our data align with this perspective: despite increased JUN expression, there is impaired activation of downstream target genes, suggesting a maladaptive injury response in CMT1A. To date, mechanistic studies involving RUNX2 knockdown in CMT1A are lacking. However, RUNX2 loss- and gain-of-function models in sciatic nerve injury provide valuable insights into its physiological role [27], [67], [71]. Intriguingly, recent findings show that depletion of RUNX2^+^ SCs, whose number increases with aging, partially restores regenerative capacity after nerve injury, implying a detrimental role for this population, in contrast to the beneficial effects associated with JUN^+^ cells [72].

Another aspect emerging from our analyses is the potential involvement of endoneurial fibroblasts in CMT1A pathology. Notably, SMOC2 expression was detected selectively in endoneurial fibroblasts but not in SCs, and its robust upregulation in CMT1A nerves suggests that this cell population may contribute to the pathological tissue remodeling associated with the disease [73]. Although our data do not directly address the functional role of endoneurial fibroblasts, it is noteworthy that these cells share a neural crest origin with SCs and are known to proliferate in response to nerve injury [73], [74]. Furthermore, induced pluripotent stem cell (iPSC) models from CMT1A patients have revealed a differentiation shift from SC toward CD34^+^ fibroblast-like cells, indicating increased developmental plasticity in this neuropathy [75]. SMOC2 has been implicated in nociception modulation in dorsal root ganglia and in the regulation of EMT, supporting the concept of sustained and dynamic interactions among SCs, neurons, and endoneurial fibroblasts during development and pathological conditions [76], [77]. Finally, the increased expression of the calcium binding protein SMOC2 in CMT1A aligns with the enrichment in GO terms associated with calcium ion sequestration pathways identified in our dataset, further suggesting that extracellular matrix and ionic homeostasis alterations may contribute to disease-associated nerve remodeling.

In conclusion, we demonstrate persistent cellular proliferation in CMT1A nerves, at least in part mediated by increased RUNX2 expression within a multicellular context that contributes to extensive tissue remodeling. These findings identify potential therapeutic avenues, including targeted inhibition of proliferation, modulation of RUNX2 activity, and promotion of lineage-specific transcriptional reprogramming within CMT1A nerve tissue. Certainly, further studies are needed to elucidate this complex and defective multicellular mechanism, which is evident in both the development and maintenance of the peripheral nervous system in CMT1A.

## Supporting information

Uncropped western blots

## Data availability statement

The authors confirm that the data supporting the findings of this study are available within the article and its supplementary materials. The RNA-seq dataset are available upon request.

## Credit authorship contribution statement

**Davide Visigalli:** Conceptualization, Investigation, Writing - original draft. **Mattia Camera:** Investigation, Visualization, Writing - original draft. **Davide Ceresa:** Data Curation, Visualization, Writing - review and editing. **Giovanna Capodivento:** Investigation, Funding acquisition. **Daniela Marrubbi:** Investigation. **Angelo Schenone:** Project administration. **Paolo Malatesta:** Supervision. **Lucilla Nobbio:** Conceptualization, Supervision, Funding acquisition, Writing - review and editing.

