## Supplementary figures for "Chronic Proliferation and Glia Heterogeneity in CMT1A: Involvement of RUNX2-Positive Schwann Cells and Endoneurial Fibroblasts"

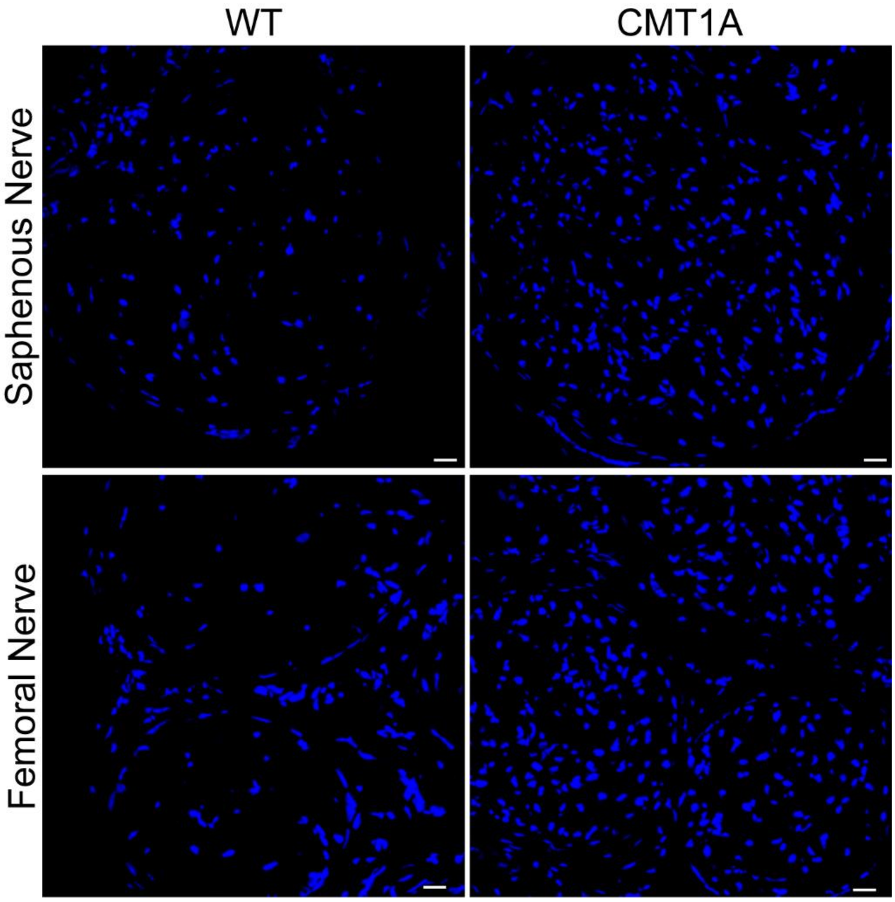


**Supplementary Figure 1. Hypercellularity in Other Peripheral Nerves.**
Representative sections of the saphenous (predominantly motor) and femoral (predominantly sensory) nerves from 12-month-old WT and CMT1A rats, stained with DAPI. Scale bars: 20 **μ**m

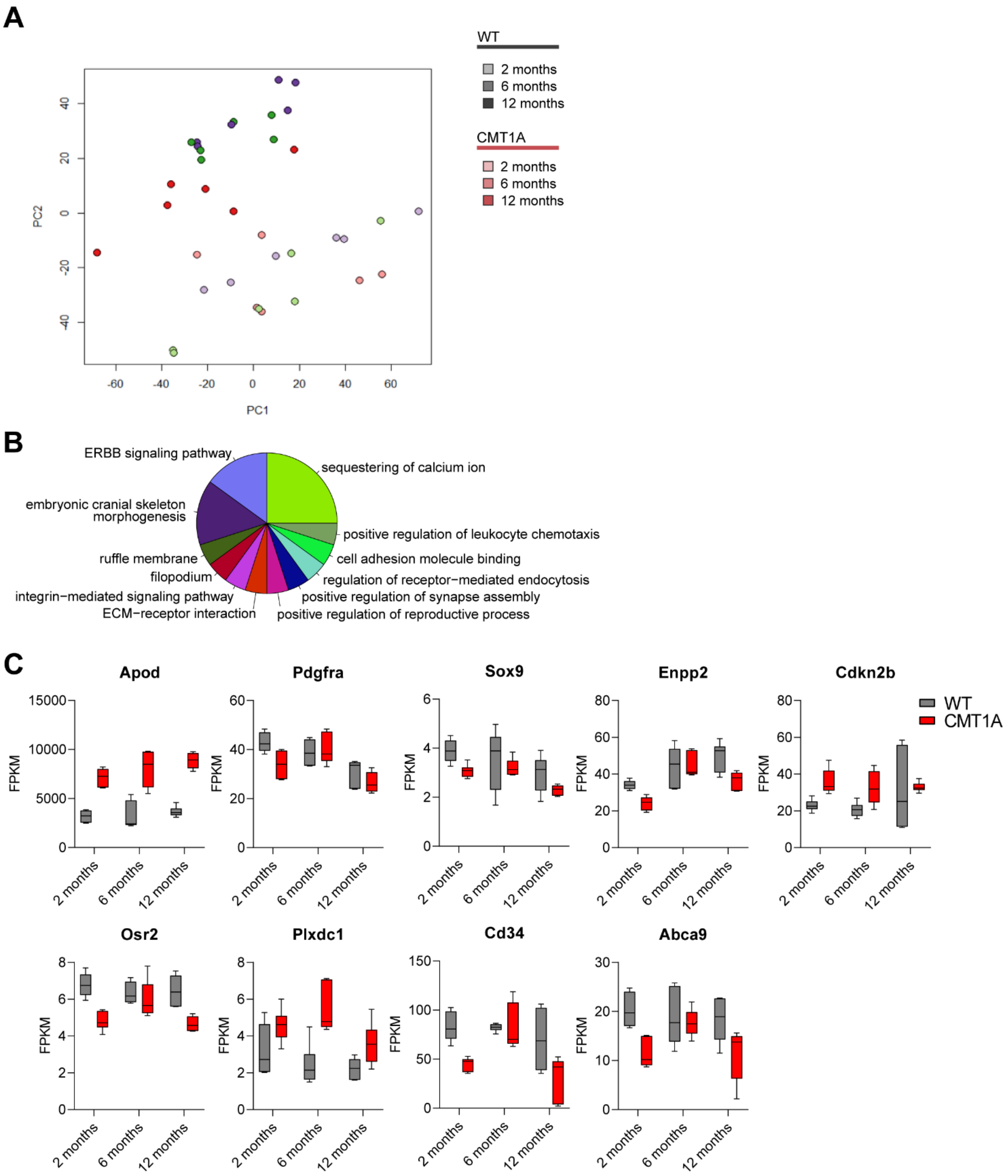


**Supplementary Figure 2. PCA and Gene Ontology Analyses.**

(A) Unsupervised principal component analysis of bulk RNA-seq profiles from WT and CMT1A sciatic nerves collected at 2, 6, and 12 months of age. (B) GO term Enrichment analysis of transcripts upregulated in 12-month-old CMT1A nerves compared with WT controls. Functionally related terms and pathways were grouped based on their shared genes. Each sector represents a functional group, with sector size indicating the proportion of enriched terms assigned to that group. The label indicates the most representative term within each group. (C) Expression levels of established markers of endoneurial fibroblasts in WT and CMT1A nerves.
