## Supplementary figures and images for "Chronic Proliferation and Glia Heterogeneity in CMT1A: Involvement of RUNX2-Positive Schwann Cells and Endoneurial Fibroblasts"

### Uncropped western blots

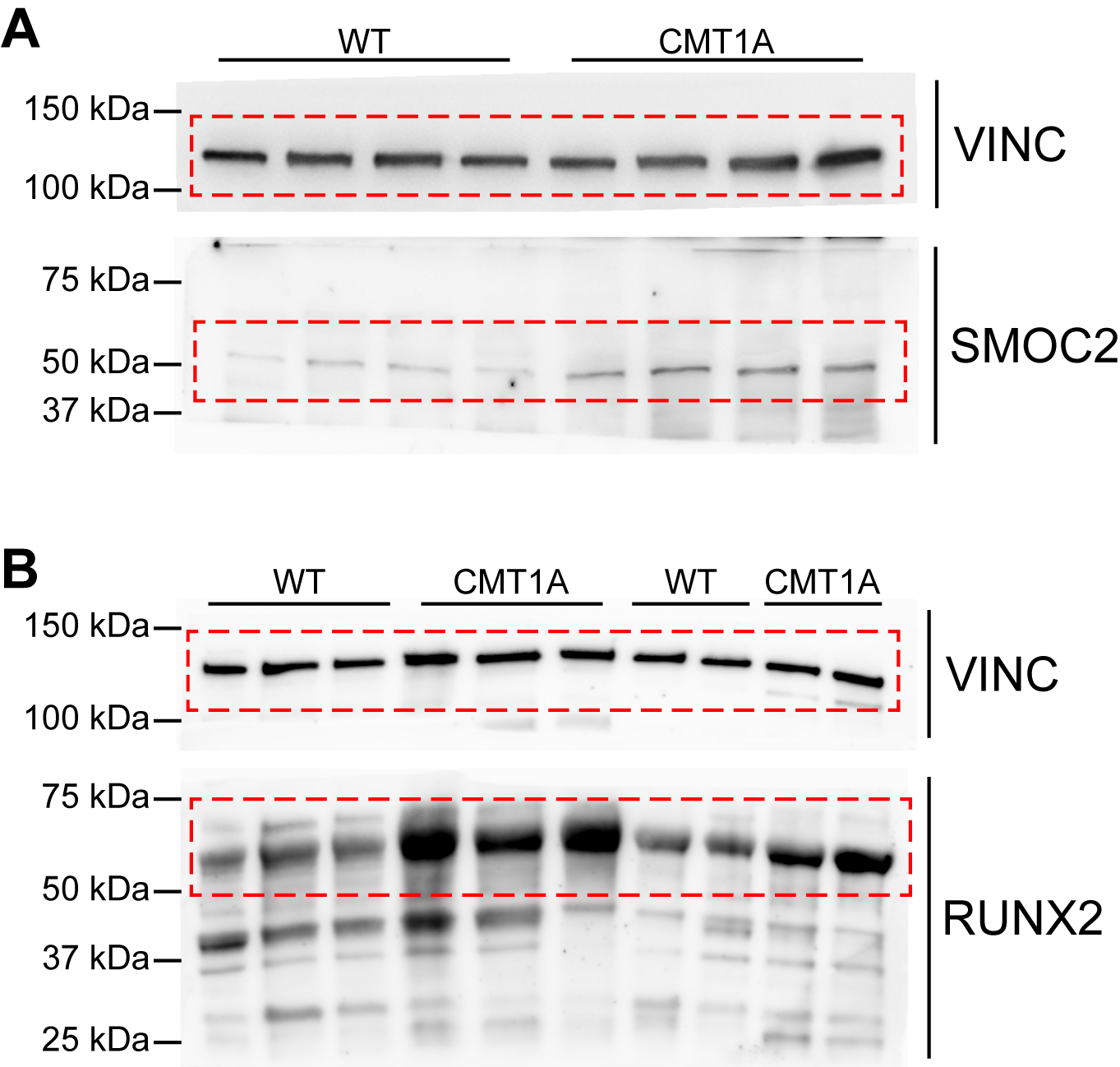
